# Fetal neural progenitors process TLR signals from bacterial components to enhance proliferation and rework brain development

**DOI:** 10.1101/2021.10.19.464985

**Authors:** Beth Mann, Jeremy Chase Crawford, Kavya Reddy, Josi Lott, Yong Ha Youn, Geli Gao, Cliff Guy, Ching-Heng Chou, Daniel Darnell, Sanchit Trivedi, Perrine Bomme, Allister J. Loughran, Paul G. Thomas, Young-Goo Han, Elaine I. Tuomanen

## Abstract

Bacterial cell wall, a universal pathogen-associated molecular pattern (PAMP), crosses the placenta into the fetal brain. We determined that PAMPs interact with TLR2/6 on murine fetal neural progenitor cells (NPCs) to induce overexpansion of all neocortical layers leading to a larger, folded cortex and abnormal postnatal behavior. The NPC overexpansion originated at E10 and targeted ventricular radial glia (vRG), the primary NPC, by shortening cell cycle and increasing self-renewal. The mechanism involved two novel signaling pathways in NPCs mediated by recognition of bacterial PAMPs by TLR2/6 including: a) loss of primary cilia, activation of hedgehog signaling, and increased FOXG1 and b) increased PI3K/AKT activity. These findings reveal PAMP/TLR2/6 acts as a morphogen in fetal neurodevelopment. In addition, the loss of *Tlr2* or *Tlr6* without pathogenic challenge, increased the number of neurons, establishing the requirement for an endogenous TLR2 signal for normal neurodevelopment in the embryo.

## Introduction

All bacteria take their shape from the highly conserved cell wall, a polymerized carbohydrate and amino acid network (Pasquina-Lemonche et al., 2020) that is recognized by elements of the innate immune response as a pathogen-associated molecular pattern (PAMP; Wolf and Underhill, 2018). Subcomponents of this complex macromolecule incite signaling by innate immune Toll-like receptor 2 (TLR2) to stimulate inflammation during acute infection (Yoshimura et al., 1999). Studies of the interactions of bacterial cell wall (BCW) with host cells in the central nervous system have focused on bacterial meningitis, a devastating infection with high mortality and severe sequelae in survivors. The molecular vocabulary encoded in the cell wall presents a diverse library that evokes all the signs and symptoms of meningitis (Tuomanen et al., 1985; Tuomanen et al., 1985). Within this inflammatory milieu driven by TLR2 signaling, neuronal injury arises by many mechanisms, including apoptosis, necroptosis, and necrosis (Braun et al., 1999; Braun et al., 2001; Aliprantis et al., 1999; Braun et al., 2002; Mitchell et al., 2004; Orihuela et al., 2006). Attenuation of neuronal cell death during human meningitis is a goal of adjunctive anti-inflammatory therapy (Tuomanen, 1990; Odio et al., 1991), but the complexity of death pathways makes directed interventions challenging in the clinic. One possible mitigation strategy to consider would be investigation of proliferation and differentiation of neural progenitor cells (NPCs) as a source of new neurons.

Although well known to drive inflammation during human infection, BCW components are also released from the gut microbiome and circulate in serum at a continuous low level, and direct non-inflammatory host responses (Clarke et al., 2010; Cryan and Dinan, 2012; Hsiao et al., 2013; Huang et al., 2019; Gonzalez-Santana and Diaz Heijtz, 2020). For instance, such chronic, tonic signals are believed to contribute to neurodegeneration and behavioral abnormalities (Laman et al., 2019; Rolls et al., 2007; Arentsen et al., 2017). This suggests that BCW components have a range of effects on the brain, the breadth of which is incompletely understood. Moreover, we know little about how bacterial infection and antibacterial treatments during pregnancy affect fetal brain development despite ample evidence that prenatal bacterial infections alter brain development and increase risks for neuropsychiatric disorders (al-Haddad et al., 2019; Christensen et al., 2019; Rakoff-Nahoum, 2016; Lowe et al., 2008; Sorenson et al, 2008; Hornig et al., 2018). Along these lines, TLR2 is present even in the early stages of development of the mammalian brain (Okun et al., 2010; Kaul et al., 2012). We have shown that BCW purified and injected into mothers crosses the murine placenta, accumulates in the fetal brain, induces abnormal expansion of brain architecture, and correlates with abnormal postnatal behavior (Humann et al., 2016). This process involved strong induction of TLR2 signaling. Thus, TLR2 ligands induce different events in the pre- and post-natal mammalian central nervous system that range from neuronal gain to neuronal loss and, at or near birth, there is a switch in outcomes driven by innate immunity from NPC proliferation to neuron death.

The primary NPCs are ventricular radial glia (vRGs; also called apical radial glia) that reside in the ventricular zone (VZ) lining the brain ventricle, each projecting a long process to the brain surface guiding migration of new excitatory neurons (Fig 1A; Kriegstein and Alvalrez-Buylla, 2009; Florio and Huttner, 2014). vRGs can divide symmetrically to increase their pool or asymmetrically to produce one vRG (preserving their pool) and one progeny that differentiates into a neuron, an intermediate progenitor cell (IPC), or an outer radial glia (oRG; also called basal radial glia). IPCs and oRGs form the subventricular zone (SVZ), where they divide to produce neurons (Hansen et al., 2010; Fietz et al., 2011). Young neurons migrate along the radial processes of vRGs and oRGs to the cortical plate, where they mature and form the 6-layered neocortex in an inside-out manner: late-born neurons migrate past early-born neurons to form a more superficial layer closer to the brain surface. Therefore, NPCs control the size and architecture of brains as both the origin and guide of new neurons. Understanding the mechanism regulating NPCs is crucial to understanding brain development and evolution and neuropsychiatric disorders caused by genetic and environmental factors.

**Figure 1.**
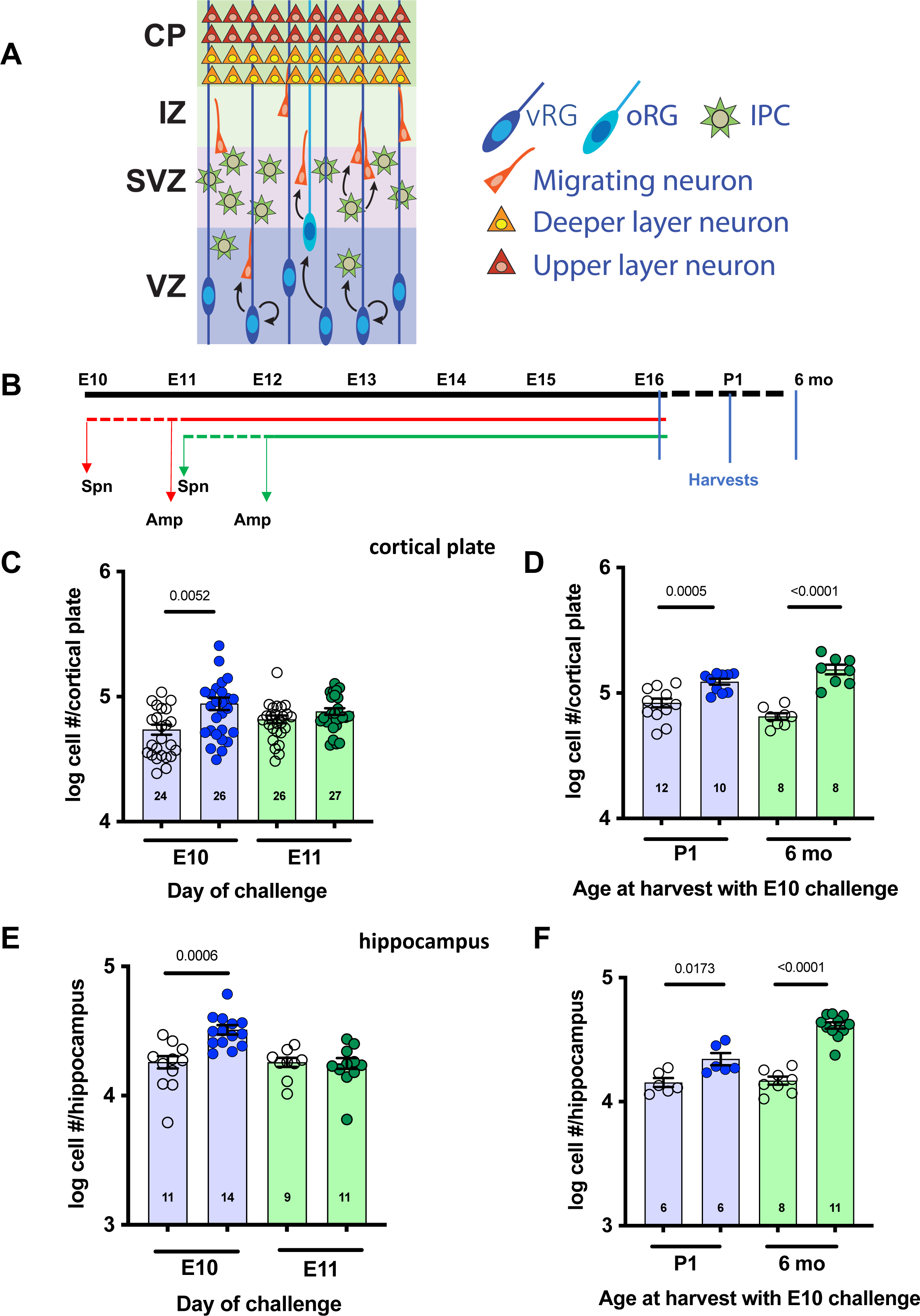
Determining the window of vulnerability for aberrant increase of neurons. **A**) Schematic representation of developing cortex with NPCs (vRGs, oRGs, and IPCs) in the ventricular zone (VZ) and subventricular zone (SVZ), migrating neurons in the intermediate zone (IZ), and differentiated neurons in the cortical plate (CP). **B)** Diagram of experimental design. Groups of dams (n>3) were challenged intratracheally with S. *pneumoniae* (Spn) T4X bacteria at 1x10^6^ cfu or PBS (control) at E10 (red dashed line) or E11 (green dashed line). 24 h later ampicillin (Amp) was administered intraperitoneally twice daily until E16 in all groups (solid lines). Brains were harvested at E16, postnatal day 1 (P1), or 6 months of age as indicated. **C-F)** Total number of neurons in the cortical plate (C,D) or the hippocampus (E,F) obtained from Nissl stained sections analysed by stereology. Measurements were taken of both right and left sides of the cortex for 3-5 dams with 2-8 embryos/dam for each experimental group: Amp alone control group (**O**) vs Spn + Amp group (•). The average of 3 sections per embryo was calculated (each symbol). *P*-values were determined by unpaired, two-tailed t-tests.

To investigate the mechanism by which bacterial debris released during treatment of maternal infection impacts fetal brain development and its potential clinical relevance, we established a model to challenge pregnant mice with live pneumococci followed by treatment with a bacteriolytic antibiotic that would flood the fetus with BCW debris while curing the mother. This model approximates the clinical scenario of the treatment of pneumococcal pneumonia in early pregnancy. We describe that bacterial lytic debris, in a restricted window of vulnerability, targeted vRGs to enhance proliferative potential, resulting in an expanded NPC pool that subsequently spread to increase numbers of neurons in the cortex with lifelong impacts on brain morphology and behavior. A greater number of founding vRGs was achieved by increased proliferation and greater self-renewal at early stages of neurogenesis. Expanded vRGs yielded more IPCs and oRGs, leading to supernumerary neurons in all layers of the neocortex and emergence of the cortical folding characteristic of brains of higher mammals but normally absent in mice. We show that this reprogramming of cortical development was driven by TLR2/6 signaling that affects PI3K/AKT signaling, primary cilia, and hedgehog (HH) signaling. Notably, the loss of either *Tlr2* or *Tlr6* increased the number of neurons in the absence of bacterial challenge, indicating that an endogenous TLR signal impacts normal neurodevelopment in the embryo. Taken together, these events define a non-canonical TLR2/6 signaling axis directly involved in neurodevelopment that reshapes brain architecture and behavior.

## Results

### A restricted developmental window vulnerable to abnormal cerebral expansion

E10 marks the initiation of neuron production and migration to form cortical neuronal layers (Kriegstein and Alvalrez-Buylla, 2009; Florio and Huttner, 2014; Homem et al., 2015). We previously showed that injecting mothers with purified BCW at E10, but not E15, increases the number of cortical neurons at E16 (Humann et al., 2016). We sought to precisely define the boundaries of the window of vulnerability and the kinetics of this abnormal event in neurogenesis and extend the impact of the model to the clinical scenario of production of BCW by treatment of pneumonia. Groups of mothers were challenged with living *Streptococcus pneumoniae TIGR4X* (Spn) at E10 or E11 to cause pneumonia without fetal infection. One day later, we injected ampicillin (Amp) to generate a rapid release of BCW components in the maternal blood stream as bacteria underwent lysis and the maternal infection was cured. We harvested fetal brains at E16, postnatal day 1 (P1), and 6 months after birth to assess cortical architecture (Fig 1B). The number of neurons significantly increased in brains of E16 embryos from dams challenged at E10, but not E11, compared to the uninfected Amp-treated control group (Fig 1C). This was also true for cortical size after E10, but not E11, challenge [E10: control 1.8e6±0.1e6 um^2^ (n=28); Spn 2.2e6±0.1e6 um^2^ (n=32), *P* = 0.0055; E11control 1.5e6±0.2e6 um^2^ (n=43); Spn 1.3e6±0.2e6 um^2^ (n=43), *P* = 0.49]. These changes persisted at P1 and 6 months of age in the E10 challenge group (Fig 1D). Cell numbers also increased in the hippocampus after challenge at E10, but not E11 (Fig 1E, F). These results identified a discrete time in neurogenesis when response of the murine fetal brain to BCW components is associated with an abnormal increase of neurons in two major areas of the brain that persisted postnatally.

It has been suggested that fetal exposure to microbial products from maternal infection is linked to permanent disorders of behavior or neuropsychiatric diseases (al-Haddad et al., 2019; Christensen et al., 2019; Rakoff-Nahoum, 2016; Lowe et al., 2008; Sorenson et al, 2008; Hornig et al., 2018). In our previous work, pups exposed *in utero* to purified BCW showed subsequent postnatal cognitive deficits (Humann et al., 2016), revealing that BCW is at least one such bioactive microbial product. We performed behavioral assays to determine if, in a live pneumococcal infection model where treatment recapitulates standard clinical care, the altered brain architecture found up to 6 months after birth was also associated with abnormal behavior (Fig 2). In a spatial recognition assay at 6 months of age, bacteria-challenged mice were less likely to enter a novel arm of the Y-maze, while control mice explored both original and novel arms equally. We tested excessive grooming, indicative of repetitive behavior, and social interactions. In the control group, mice significantly decreased the amount of time they spent self-grooming with age, while all the bacteria-challenged mice demonstrated high levels of repetitive behavior. In a sociability assay at 6 months of age, the challenged group interacted significantly less with a companion mouse than the control group did. These results indicated that a treated maternal infection, which released BCW that crossed the placenta early in pregnancy, resulted not only in abnormal brain architecture but also disordered spatial memory, excessive repetitive behavior, and antisocial behavior.

**Figure 2.**
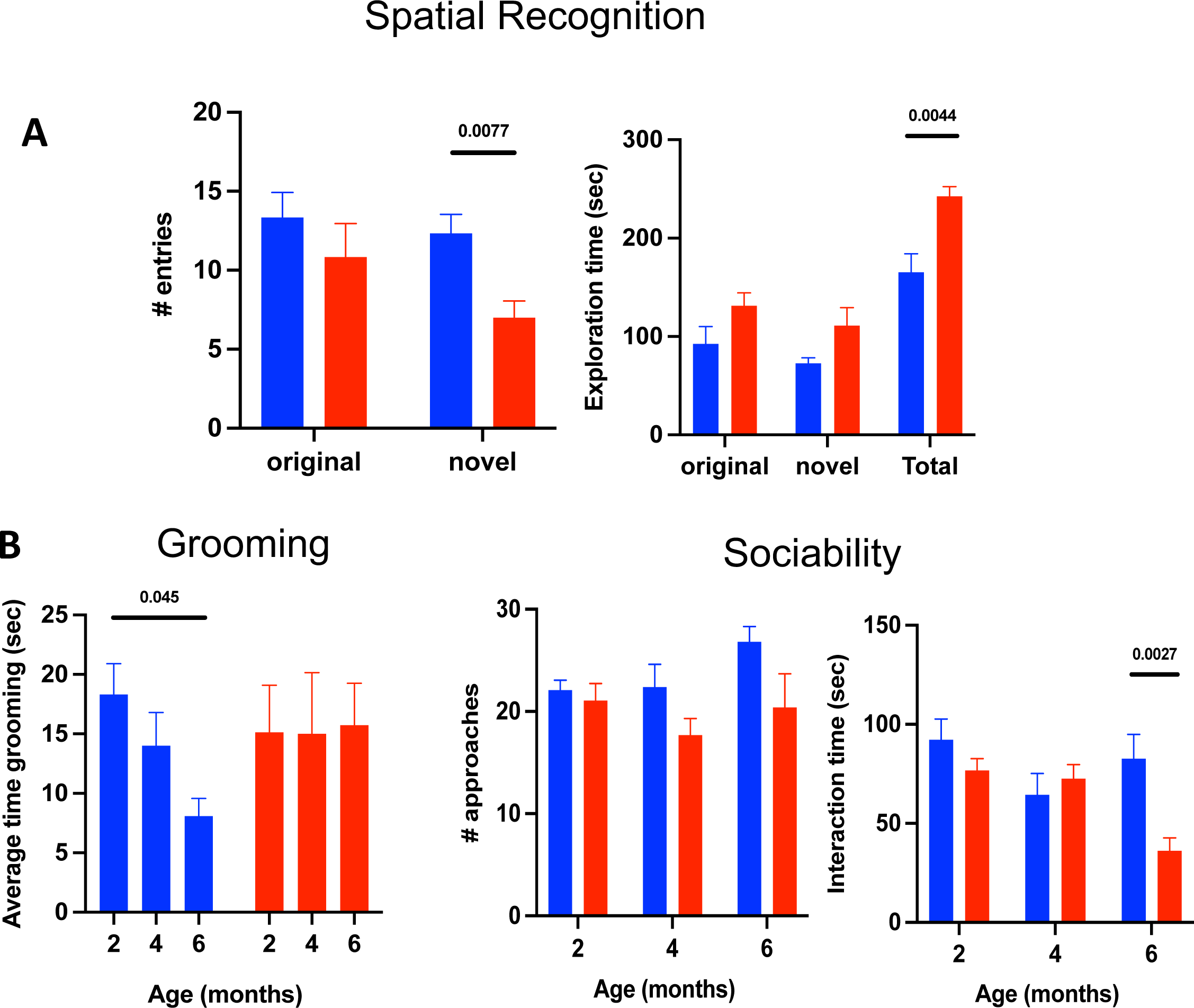
Behavioral consequences of aberrant increase of neurons. **A)** Spatial recognition was tested in a Y maze comparing 6-month-old mice exposed in utero at E10 to either Amp alone (blue bars) or Spn + Amp (red bars) for number of entries into a novel arm. **B)** Pups were tested at 2, 4, and 6 months of age in two assays. Repetitive behavior was assessed by time spent grooming. Sociability was assessed in a white box as the time spent at a transparent, perforated partition upon addition of a non-littermate mouse. Values represent mean ± standard deviation of 2-3 assays, 4-6 mice per assay. *P*-values were determined by unpaired, two-tailed t-tests.

### Neurons increase in all cortical layers

We sought to define which layers of excitatory neurons were increased by BCW challenge. Neurons in different cortical layers were marked by specific transcription factors: TBR1 for layer VI, which arises early in development; CTIP2 (also known as BCL11B) for layer V, which arises next; and SATB2 for layers II-IV, which arise last in development (Arlotta et al., 2005; Britanova et al., 2008; Hevner et al., 2001). Bacterial challenge did not affect the cortical layering: VI innermost and II outermost (Fig 3A). However, the E10 challenge, but not E11, significantly increased the number of neurons in each layer (Fig 3B) even though most SATB2+ neurons of layers II-IV arise at E14-E17 and most TBR1+ layer VI neurons arise at E11.5-E13. Since BCW is cleared from the fetal brain within 2-3 days (Humann et al., 2016), these findings suggest a persistent dysregulation of neurogenesis long after exposure to BCW.

**Figure 3.**
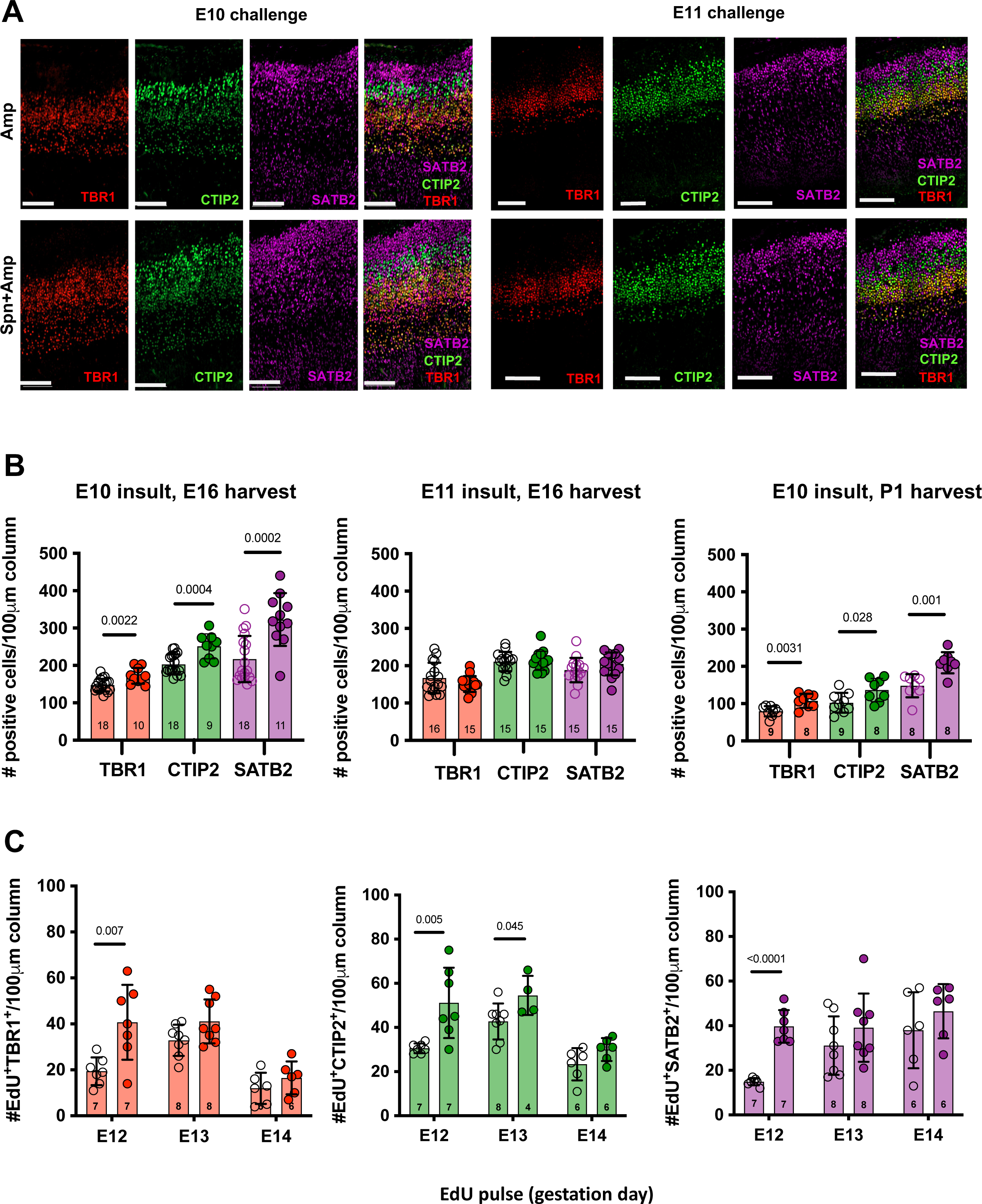
PAMP challenge increases number of all cortical excitatory neurons. Dams were challenged as shown in Fig. 1B and then harvested at E16 or P1. **A)** Neuronal layers were stained for TBR1 (red, layer VI), CTIP2 (green, layer V), or SATB2 (purple, layer II-IV). The scale bar represents 100 µm. **B)** Graph shows quantification of neurons in each layer in a 100 µm cortical column from the base of the VZ to the top of the cortical plate: Amp alone control group (**O**) vs Spn + Amp (•). Measurements were taken from at least 3 dams per condition with 2-7 embryos/dam. Each symbol averages 2 sections from one embryo, and each bar represents the mean ± SEM. **C)** To follow the production of each of 3 cortical neuron types, EdU was given on E13, E14, or E15, and embryos were harvested at E16. Graph shows quantification of EdU-positive cells co-labeled for each neuronal marker in a 100 µm cortical column: Amp alone (○) vs Spn + Amp (•). Each condition was tested in 2-3 embryos from 4 independent experiments. Data are shown as mean ± SEM. *P*-values were determined by unpaired two-tailed t-tests.

To examine the kinetics of neuronal increase in the E10 challenge model, newborn cells were labeled by a pulse of EdU at E12, E13, or E14 and identified at E16 (Fig 3C). While the challenged group exhibited more EdU+ cells than the control group at all time points for all neuron types, the peak number of EdU+ neurons for each type occurred in the order in which they arise during normal development: TBR1 (layer VI) was highest at E12, CTIP2 (V) at E13, and SATB2 (II-IV) at E14. Notably, EdU injected at E12 strongly labeled TBR1+ cells but weakly labeled SATB2+ cells (Supplementary Fig S1), indicating that TBR1+ cells were produced from an NPC division directly following EdU incorporation at E12, whereas SATB2+ cells were produced after multiple rounds of NPC divisions that diluted incorporated EdU. These results indicate that more NPCs that incorporated EdU at E12 expanded and self-renewed in the challenged group than in the control group, resulting in an expanded pool of NPCs producing even late-born SATB2+ neurons.

### Rapid initial expansion of vRGs by changes in cell cycle time

vRGs are the primary NPCs that produce neurons, either directly or through oRGs and IPCs (Fig 1A). We hypothesized that an initial expansion of vRGs followed by increased production of oRGs and IPCs could create a progressive wave of increased neurons in each layer of the cortical plate. Therefore, to test if E10 challenge, but not E11, caused initial expansion of vRGs, we quantified vRGs one day after Amp treatment had initiated the release of BCW. Consistent with our hypothesis, vRGs (PAX6+ TBR2-cells in the VZ) but not IPCs (TBR2+ cells) were greatly expanded after E10, but not E11, challenge (Fig 4 A, B), indicating that vRGs amplified themselves as primary targets of BCW.

**Figure 4.**
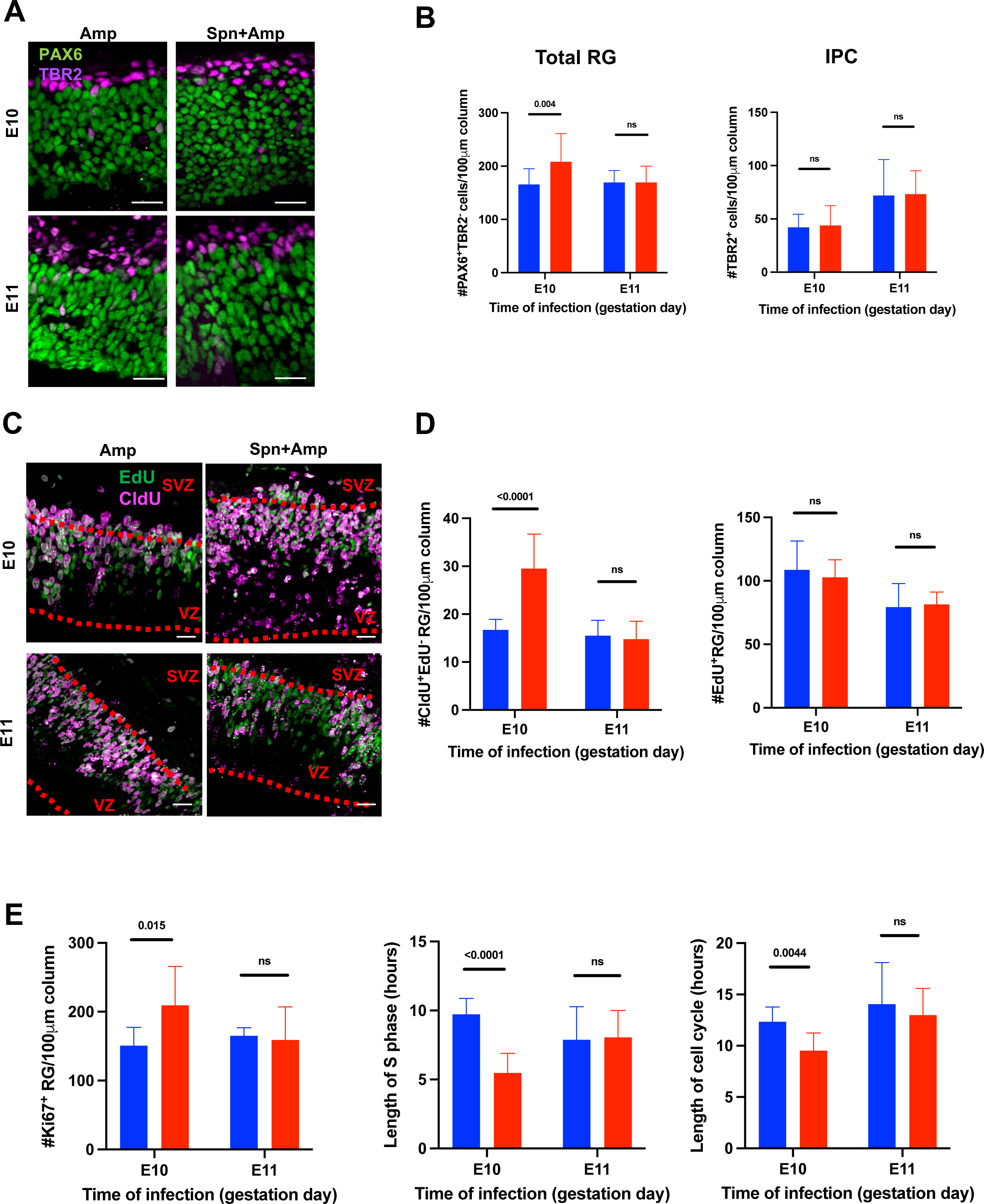
PAMP challenge shortens vRG cell cycle. **A, B)** We challenged dams at E10 or E11 (Blue: PBS, Red: Spn), treated them with Amp 24 h later, and NPCs were quantified RGs (PAX6+TBR2-cells) and IPCs (PAX6-TBR2+) at 48 h (E12 or E13, respectively). Total RGs were counted as PAX6+TBR2-cells, and IPCs were counted as PAX6-TBR2+. **C-E)** To determine cell cycle kinetics of RGs in the 48 h after challenge at E10 or E11 (as in panel A; Blue: Amp, Red: Spn + Amp), double thymidine analogue labeling was carried out by successive injections of CldU at 2 h before harvest and EdU at 0.5 h before harvest. Note the many CldU+EdU-cells in the lower VZ in embryos that were challenged with bacteria at E10. Graphs show RGs in S phase (EdU+), RGs that had left S phase (CldU+EdU-), RGs in cell cycle (Ki67+), and the lengths of S phase and the cell cycle of RGs. Blue: Amp, Red: Spn + Amp. Each condition was tested in 10-12 embryos from 3 independent experiments. Data are shown as mean ± SEM. *P*-values were determined by unpaired two-tailed t-tests.

We next investigated how bacterial challenge expanded vRGs. Cell-cycle kinetics is closely associated with the division mode of NPCs (Dehay and Kennedy, 2007). In particular, vRGs undergoing proliferative self-amplifying divisions show a shorter cell cycle than vRGs producing differentiating progenies (Calegari et al., 2005). Thus, we investigated if the expanded vRGs were characterized by a shortened cell cycle by using a double thymidine analogue labeling method with successive injections of CldU at 2 hours (h) and EdU at 0.5 h before harvest (Nowakowski et al., 1999; Hayes and Nowakowski, 2000). EdU+ PAX6+ TBR2-cells represent vRGs in S phase at the time of harvest, whereas CldU+ EdU-cells represent vRGs that have left S phase and entered G2 phase during the 1.5 h between CldU and EdU injections (Fig 4C). The nuclei of vRGs show interkinetic migration (Miyata et al., 2014): nuclei undergo S phase at the upper part of the VZ, move down to the ventricular surface during G2 phase, and divide at the ventricular surface. Consistent with interkinetic nuclear migration of vRGs, the nuclei of vRGs in S phase (EdU+) were concentrated in the upper part of the VZ (Fig 4C). Remarkably, many CldU+ EdU-vRG nuclei were present in the lower VZ in BCW-challenged but not control E10 embryos, suggesting that more cells exited S phase and entered G2 phase during the 1.5 h interval in E10-challenged than control embryos (Fig 4C). Accordingly, E10 challenge dramatically increased the number of CldU+ EdU-cells compared to control embryos (Fig 4D). Despite the increased exit from S phase, the number of vRGs in S phase (EdU+ cells) did not change, indicating that the number of vRGs entering S phase was also increased (Fig 4D). These results suggested a shortened S phase and cell cycle. Indeed, E10, but not by E11, challenge significantly shortened the lengths of S phase and the cell cycle (Fig 4E) calculated according to Nowakowski et al. (1989). Together, these results show that E10, but not E11, bacterial challenge, expanded vRGs within one day after Amp treatment by shortening the cell cycle length.

### Initial expansion of vRGs leads to expanded NPCs in later stages and cortical folding

The increase of all cortical layers of neurons (Fig 3B) suggests that the early expansion of vRGs by two days after E10 challenge (Fig 4B) was maintained and propagated into downstream cell lineages throughout the neurogenic period. To test this possibility, we quantified the number of vRGs, oRGs, and IPCs at E16, a late stage of cortical neurogenesis. All 3 cell types were increased solely in the E10 challenged group, consistent with a time-restricted vulnerability of vRGs to bacterial PAMPs (Fig 5A, B). The increase of oRGs and IPCs indicates that the initial expansion of vRGs resulted in subsequent expansion of downstream NPCs. Consistent with this, new IPCs with residual vRG markers (TBR2+ PAX6+) were nearly doubled in the VZ in the challenged group (mean ± SD of 11.8 ± 6 for Amp control and 19.88 ± 6 for challenged group, p=0.042). These results indicated that E10 challenge expanded all NPC types by E16.

**Figure 5.**
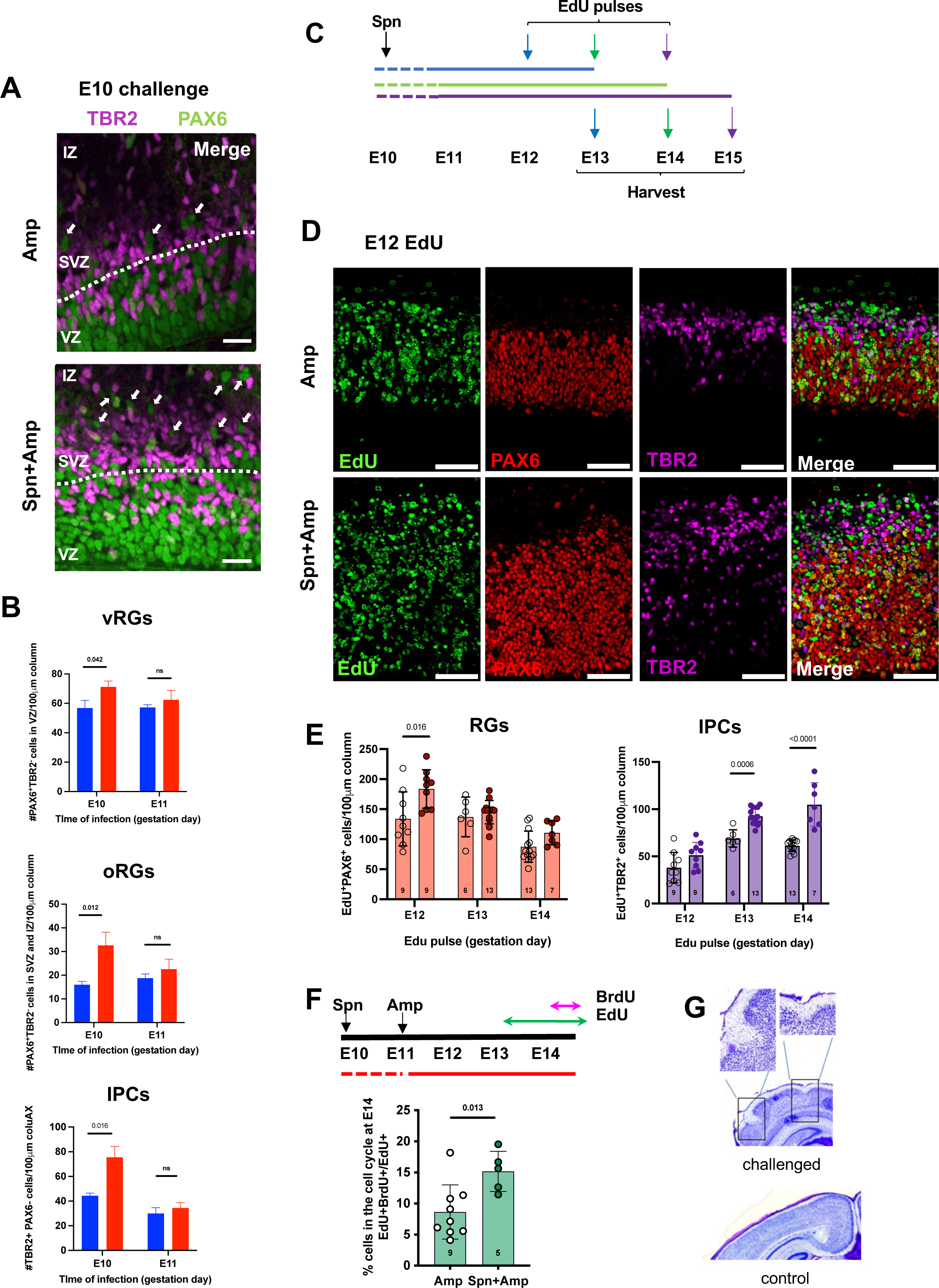
Propagation of expanded vRGs into oRGs and IPCs. **A, B)** We challenged dams with bacteria challenge at E10, and quantified NPCs at E16 by both PAX6 and TBR2 markers as well as location: vRGs in the VZ, oRGs outside the VZ, and IPCs in the SVZ, regions defined by white dotted line (Blue: Amp, Red: Spn + Amp). **C - E)** To follow expanded NPCs in later developmental stages, we gave pulses of EdU were given at E12, E13, or E14, and quantified NPCs in the VZ and SVZ at harvest 24 h later (scheme panel C). The numbers of EdU+ RGs and IPCs were determined in challenged (•) vs controls (○). The scale bar represents 100 µm. **F)** NPC self-renewal was assessed by labeling with an EdU pulse at E13 (green arrow) followed in 24 h by a BrdU pulse (pink arrow) 1.5 h before harvest. Self-renewed cells (EdU+BrdU+) were counted and expressed as a % of total EdU+ cells. Graph represents the mean ± SEM of 3 sections per embryo and 2-5 embryos from 2 independent experiments. *P*-value was determined by unpaired two-tailed t-test. **G**) Nissl staining at P10 shows cortical folding in E10 challenged (top) but not in control (bottom) embryos. Boxes indicate cortical fold.

To understand how expanded vRGs propagate to expanded NPCs in later developmental stages, we labeled proliferating NPCs by pulses of EdU at E12, E13, or E14, and cells in the VZ and SVZ were quantified 24 h later (Fig 5C-D) (Franco et al., 2012; Ponti et al., 2013). EdU injection labeled more RGs at all ages in the challenged group than in control, with the highest increase by E12 injection, while EdU+ IPCs increased gradually, with significant increases at E13 and E14 injection times. These results suggest that more vRGs expanded at E12 in the challenged group than in control and that increased vRGs subsequently produced more IPCs at E13 and E14. This supports a model that E10 challenge resulted in a wave of expansion of both RGs and IPCs over time in their expected developmental sequence, with vRGs impacted first, followed by IPCs and oRGs.

Next, we determined whether the expanded NPCs were sustained into later stages through self-renewal. We injected dams with EdU at E13 and with BrdU at 1.5 h before harvest at E14 (Fig 5F) to identify progenitors that proliferated at E13 (EdU+), remained as progenitors, and proliferated again at E14 (BrdU+). The number of self-renewed progenitors (EdU+ BrdU+) was significantly increased in E10 challenged group. These findings showed that the initial expansion of vRGs at E12 propagated to expansion of all 3 NPC types at later stages in E10 challenged embryos through increased proliferative capacity and self-renewal.

Initial expansion of vRGs eventually yielded more IPCs and oRGs, leading to the increase of neurons in all layers of the neocortex. The hallmark of this expansion is enhanced cortical folding characteristic of brains of higher mammals. Remarkably, the expansion of NPCs and cortical neurons in E10-challenged murine brains resulted in the cortical folding in 50% of challenged mice (5 of 10), a feature absent in control mice (0 of 10) (Fig 5G).

### Bacterial PAMPs require TLR2/6 to expand neurons

NPCs and embryonic neurons express TLRs that recognize bacterial PAMPs (Okun et al., 2010; Kaul et al., 2012). We previously showed that purified pneumococcal cell wall requires TLR2 to increase the number of cortical neurons (Humann et al., 2016). TLR2 is a heterodimer interacting with either TLR1 or TLR6. Classical BCW-induced inflammation in postnatal meningitis involves TLR2/1 signaling in leukocytes and microglia; any role of TLR2/6 in meningitis is unknown. Furthermore, the impact of either TLR2/1 or TLR2/6 signaling in fetal brain development is unknown. We therefore tested which TLR2 signaling pathway drove abnormal brain development following bacterial infection and treatment. We challenged pregnant mice bearing WT, *Tlr1^-/-^, Tlr2^-/-^,* or *Tlr6^-/-^* embyros with Spn at E10, treated with Amp, and examined the brains at E16. Neuronal layering was normal in all mutant embryos (Fig 6A). However, BCW challenge increased the number of cells in the cortical plate in WT and *Tlr1^-/-^* embryos but not in *Tlr2^-/-^* and *Tlr6^-/-^* embryos (Fig 6A, Supplemental Fig S2), demonstrating that the TLR2/6 axis is required for bacterial PAMPs to increase cortical neurogenesis.

**Figure 6.**
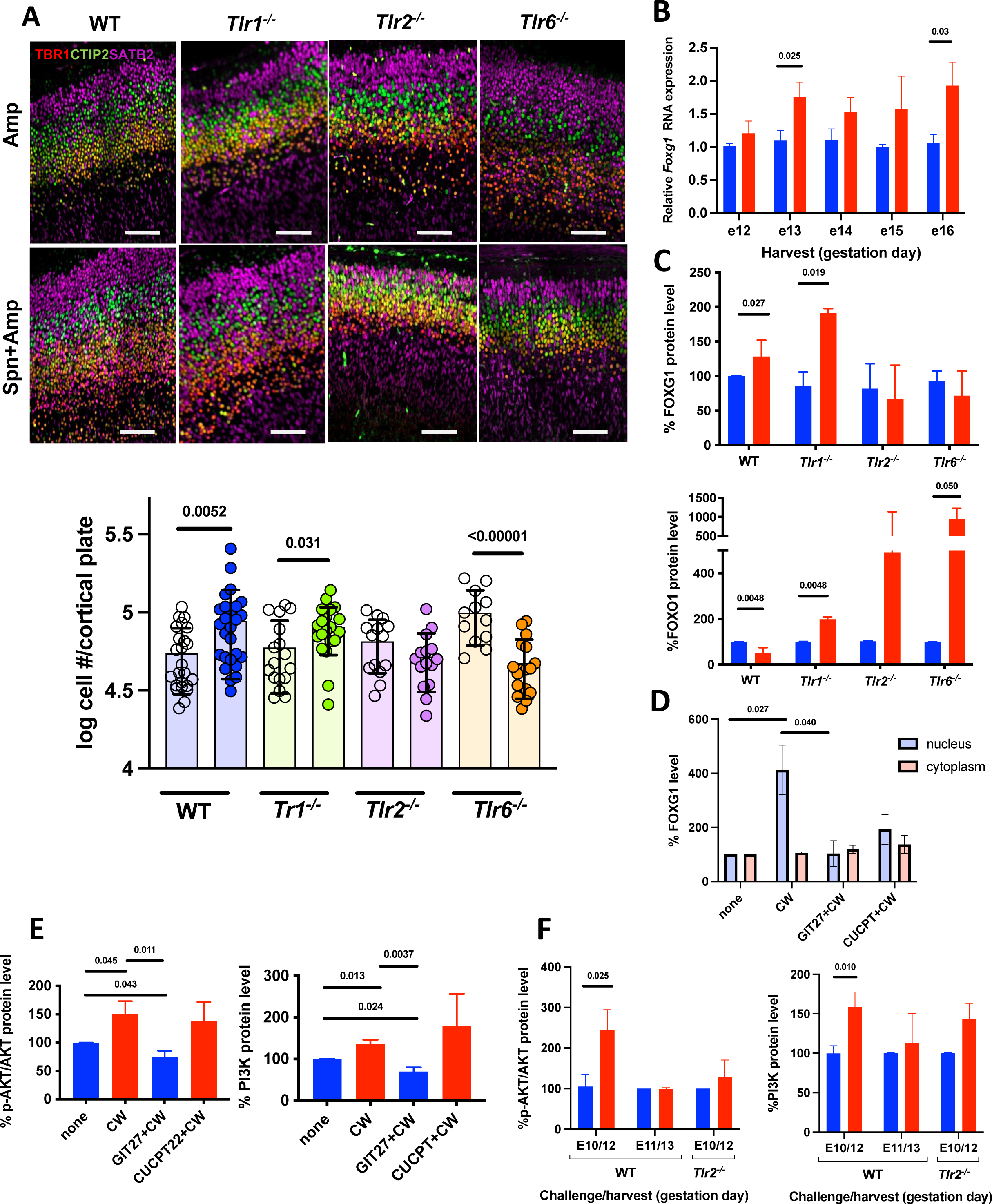
PAMPs require TLR2/6 to expand cortical neurons. **A)** Using the *in vivo* assay of Fig 1B, dams deficient in *Tlr1, 2*, or *6* were challenged with bacteria or PBS at E10. Pictures show E16 cortex stained with cortical excitatory neuronal markers. Graphs show total numbers of neurons in the cortical plate from Nissl-stained sections analyzed by stereology at E16. Right and left sides of the cortex were measured, and the average of 3 sections per embryo was calculated (each symbol). Measurements were taken for 3-4 dams with 4-8 embryos/dam for each experimental group as indicated by value inside each bar: Amp (○) vs Spn + Amp (•). Values were combined from at least 3 different experiments. The scale bar represents 100 µm. **B)** *Foxg1* transcript levels were measured by qPCR with RNA from brains exposed to Amp (blue) or Spn + Amp (red) at E10 and harvested at the indicated day. Values represent 3-4 dams and 3-5 embryo lysates/dam. Data are shown as mean ± SEM. **C)** WT and indicated *Tlr* deficient dams were challenged at E10 with PBS (blue) or Spn (red), treated with Amp 24 h later as in Fig 1B, and embryos were harvested at E16. Brain lysates were assessed by Western blot for FOXG1 and FOXO levels. Values are expressed as % of WT control set at 100, using 2-3 dams per condition with 2-4 embryos/dam. **D)** FOXG1 protein levels were measured in NPCs cultured *in vitr*o from E12.5 embryos. TLR antagonists were applied as indicated: GIT27 10ug/ml for TLR2/6 or CU-CPT22 5uM for TLR 2/1. After 2 h, cells were exposed to purified BCW (MOI 1) for 2hrs and harvested. Cell lysates were processed into nuclear and cytoplasmic fractions and assayed by Western blot for FOXG1. Protein levels are expressed as % of those in cells not treated with BCW or antagonist. Each bar represents the mean ± SEM of 4 independent experiments. **E)** pAKT and PI3K protein levels were measured in NPCs cultured from E12.5 embryos. TLR antagonists were applied as in panel D. Protein levels are expressed as % of controls not treated with BCW or antagonist. Each bar represents the mean ± SD of 4 independent experiments. **F)** We treated WT and *Tlr2-/-* dams at the indicated day with PBS (blue) or Spn (red), treated them with Amp 24 h later, and collected brains another 24 h later, as indicated. Brain lysates were assessed by Western blot for p-AKT, AKT or PI3K levels. Values are expressed as % of WT control. We used 2-3 dams per condition with 2-4 embryos/dam. *P*-values for all panels were determined by unpaired, two-tailed t-tests.

### Loss of TLR signaling affects normal brain architecture

Given the impact of TLR signaling on fetal cortical architecture in the face of bacterial challenge of the mother, we tested if loss of TLR signaling affected brain structure in <u>un</u>challenged controls. Remarkably, the number of cortical neurons significantly increased in *Tlr2^-/-^* and *Tlr6^-/-^* embryos compared with WT, whereas the number of neurons was not changed in *Tlr1^-/-^* embryos (Fig 6A). These effects in unchallenged *Tlr2^-/-^* and *Tlr6^-/-^* mice established the requirement for an endogenous TLR2 and TLR6 axis signal to set the number of neurons during normal neurodevelopment and that TLR ligands, such as bacterial PAMPs, modulate that signal. Together, our results indicate that TLR2 and TLR6 suppress cortical neuronal expansion under healthy conditions and that pathogenic TLR ligands reverse such suppression.

### TLR2/6 increases PI3K/AKT signaling and FOXG1 levels

FOXG1 is a nuclear transcriptional repressor expressed very early in development and promotes expansion of NPCs (Seoane et al., 2004; Xuan et al., 1995; Hanashima et al., 2004; Hanashima et al., 2002; Miyoshi and Fishell, 2012; Martynoga et al., 2005; Regad et al., 2007; Shen et al., 2006; Du et al., 2019). In humans, altered levels of FOXG1 is associated with a high risk of abnormal behavior and autism (Hazlett et al., 2017; al Haddad et al., 2019; Scharfman and Hen, 2007; Wong et al., 2019), a phenotype recapitulated in our mouse model. We found that maternal bacterial challenge at E10 followed by antibiotic treatment increased *Foxg1* mRNA levels in fetal brain (Fig 6B). Importantly, bacterial challenge also increased FOXG1 protein levels in WT and *Tlr1^-/-^*, but not in *Tlr2^-/-^* or *Tlr6^-/-^* embryonic brains (Fig 6C). Moreover, *in vitro* NPCs treated with purified cell wall showed a strong induction of nuclear FOXG1 that was blocked by pretreatment with TLR2/6 inhibitor GIT27 but not the TLR2/1 inhibitor CUCPT (Fig 6D), indicating that BCW signals through TLR2/6 to increase FOXG1 levels.

FOXG1 expands NPCs, at least in part by binding to and inhibiting FOXO from inducing the expression of cell cycle inhibitors (Seoane et al., 2004). PI3K/AKT signaling promotes proteasomal degradation of FOXO (Aoki et al., 2004) and cooperates with FOXG1 to inhibit FOXO (Seoane et al., 2004). Remarkably, bacterial challenge also decreased FOXO protein levels in WT embryos but not in *Tlr2^-/-^* or *Tlr6^-/-^* embryos (Fig 6C), suggesting a role for PI3K/AKT signaling downstream of TLR2/6 (Pourrajab et al., 2015). Accordingly, NPCs treated *in vitro* with purified cell wall showed a strong induction of PI3K levels and AKT phosphorylation that were blocked by pretreatment with TLR2/6 inhibitor GIT27 but not the TLR2/1 inhibitor CUCPT (Fig 6E). Moreover, pAKT and PI3K protein levels were increased in brain lysates of E10 challenged WT but not *Tlr2^-/-^* embryos (Fig 6F). Together, these data suggest that TLR2/6 signaling expanded NPCs in part via PI3K/AKT signaling and FOXG1.

### TLR2 signaling modulates ciliogenesis and HH pathway

A pathway connecting TLR2/6 signaling to FOXG1 has not been previously described. To understand further the mechanism by which TLR2/6 signaling increased FOXG1 and expanded vRGs, we used spatial transcriptomics to compare gene expression in E12 brains from dams challenged at E10 with bacteria or PBS and treated with Amp. Comparisons focused on the VZ, where vRGs reside. PI3K/AKT and HH signaling pathways were among those significantly enriched within the bacteria challenged VZs (Fig 7A, B; Supplementary Fig S3). Enrichment of PI3K/AKT signaling was in accord with the activation of PI3K/AKT signaling in the challenged group (Fig 6) and its role in promoting NPC proliferation. HH signaling induces *Foxg1* expression in NPCs (Danesin et al., 2009), thus its enrichment suggested a possible mechanism for increased *Foxg1* expression in the challenged group. Moreover, HH signaling is a conserved mechanism that induces the expansion of NPCs and the growth and the folding of the neocortex (Komada et al., 2008; Wang et al., 2016; Hou et al., 2021), all of which were the consequences of BCW challenge at E10. Notably, *Fgf15,* a target of HH signaling in the developing cortex (Wilson et al, 2012), was significantly upregulated within the E10 challenged group and remained strongly upregulated between E12 and E16 (Fig 7C). Previous work showed that loss of GLI3 repressor (GLI3R) that suppresses the expression of HH target genes in vRGs strongly increases *Fgf15* expression, which in turn shortens the cell cycle and promotes proliferation of NPCs (Wilson et al, 2012; Youn and Han, 2017). Consistent with the enrichment of HH signaling signatures and increased *Fgf15*, GLI3R levels were significantly decreased in E10 challenged fetal brains in a TLR2 dependent manner (Fig 7D).

**Figure 7.**
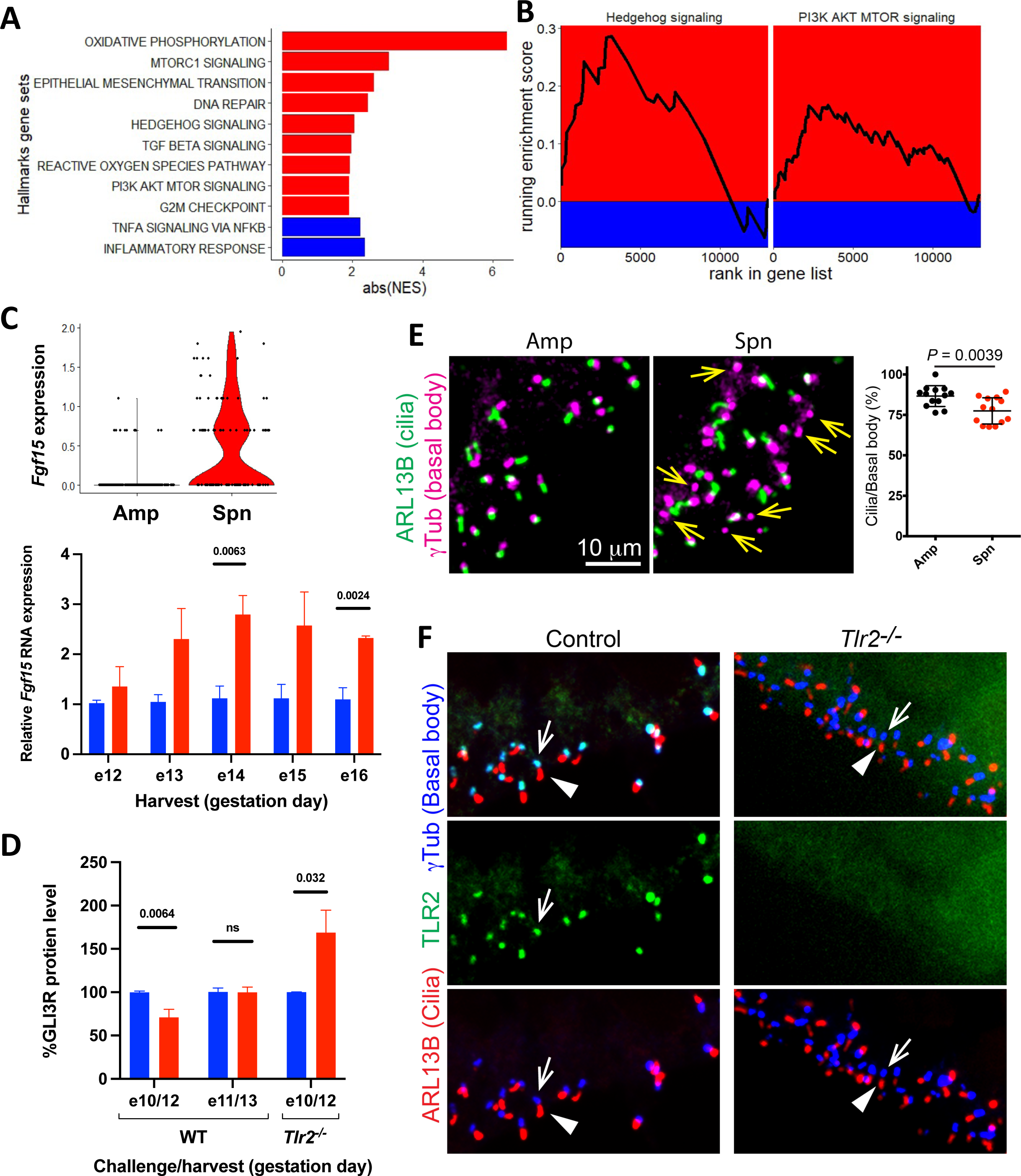
Effect of PAMP/TLR2 challenge on primary cilia and HH signaling. **(A)** Pre-ranked Gene Set Enrichment Analysis (GSEA) was performed comparing control to bacteria-challenged E12 VZ gene expression assessed by spatial transcriptomics. Absolute values of the Normalized Enrichment Scores (NES) for selected pathways of interest that met statistical significance (FDR < 0.05) are presented. Red bars correspond to pathways significantly enriched in the Spn + Amp condition, whereas blue bars correspond to pathways significantly enriched in the control (Amp) condition relative to the treated condition. **B)** Enrichment plots for two of the pathways described in (A), visualizing the running GSEA enrichment scores across ranked expression differences in each gene set. Red quadrants correspond to running enrichment scores associated with Spn + Amp brains, whereas blue quadrants correspond to those associated with control (Amp) brains. **C)** Left: Violin plot of *Fgf15* expression obtained from control (Amp) and treated (Spn + Amp) E12 VZ as assessed by spatial transcriptomics. Right: Relative expression of *Fgf15* as assessed by RT-qPCR in brain lysates of embryos exposed to Amp (blue) or Spn + Amp challenge (red) at E10 and harvested at indicated day. Values represent 3-4 dams and 3-5 embryo lysates per dam. Data are shown as mean ± SEM. *P*-values were determined by unpaired two-tailed t-test are shown above bars. **D)** Western blot analysis as per panel C for expression of GLI3R as a function of day of challenge and harvest in WT or *Tlr2^-/-^* mice. **E)** Pictures of E12 cortex stained for γTub (basal body marker, blue), ARL13B (primary cilia marker, red), and TLR2 (green). Bottom row: The basal body (blue; arrows) is located at the base of cilia (red; arrowheads). Top row (merge of middle and bottom): colocalization (cyan) of TLR2 (green, middle panel) with γTub at the base of cilia in vRGs. TLR2 staining is absent in sections from *Tlr2-/-* mice. The scale bar = 10 µm for **E** and **F**.

Next, we investigated how bacterial challenge decreased GLI3R levels. GLI3 must shuttle through primary cilia to be proteolytically cleaved to become GLI3R (Goetz and Anderson, 2010; Guemez-Gamboa et al., 2014). Accordingly, the loss of cilia in NPCs leads to the loss of GLI3R and derepression of HH target genes, including *Fgf15* (Wilson et al., 2012; Youn and Han, 2017; Hou et al., 2021; Wang et al., 2016). Remarkably, bacterial challenge at E10 significantly decreased the number of vRGs with primary cilia (Fig 7E). Moreover, TLR2 localized to the base of primary cilia in vRGs (Fig 7F). Together, these findings suggest that bacterial-TLR2 signaling at the base of cilia suppresses formation of cilia and GLI3R, leading to the derepression of HH target genes and the expansion of NPCs.

## Discussion

The maternal fetal interface monitors a constant, undercurrent dialog between the fetus and the mother, including responses to microbial products released from the maternal microbiota or by treatment of infection. Our data indicate that bacterial debris from the mother traverses the murine placenta into the fetal brain, where it directly interacts with NPCs, resulting in abnormal NPC proliferation, altered brain architecture, and aberrant postnatal behavior. The current study sought to examine this phenomenon in a context relevant to clinical maternal/fetal care where release of bacterial lytic products in the maternal bloodstream is initiated by ß lactam antibiotic therapy to cure pregnant mice with pneumococcal pneumonia. In this model, a burst of BCW release occurs over 6-12 hours after antibiotic administration as bacteria rapidly die, creating a discrete pulse of bacterial debris that accumulates in the fetal brain by 24-48 hours. Bacterial challenge at E10 increased the number of all excitatory neuron types of the neocortex and the hippocampus, indicating a wide impact of bacterial components released in the mother on fetal brain architecture. This increase in neurons resulted in an enlarged and folded neocortex, which is characteristic of the neocortex of higher mammals. These structural changes were accompanied by cognitive and behavioral abnormalities, including disordered spatial memory, excessive repetitive behavior, and antisocial behavior. These findings indicate that brief bacterial infection and treatment in the early stages of pregnancy could have long-lasting effects on fetal brain structure and function.

The bacterial challenge increased the number of excitatory neurons in all cortical layers although they arise at different times spanning E11.5 to E17.5 (Molyneaux et al., 2007). An effect initiated at E10 but spanning many days thereafter suggested that the consequences of bacterial challenge originated in the pool of NPCs and was reflected in all cells born afterward. This was confirmed, as all NPCs (vRGs, oRGs, and IPCs) underwent expansion. In particular, the greatest effect targeted the primary NPCs, the vRGs, and progressed as a wave in time through the generation of later cell types. Before producing neurons, vRGs divide symmetrically to amplify themselves. As they switch to divide asymmetrically to produce neurons, cell cycle lengthens (Calegari et al., 2005; Calegari and Huttner, 2003). Our data suggest that signals from bacterial debris expand the initial pool of vRGs at the early stage of cortical development by delaying the lengthening of their cell cycle and increasing self-amplifying divisions. This initial expansion of vRGs was propagated to the expanded pool of IPCs and oRGs that were also maintained by increased self-renewal. It is notable that bacterial debris expanded both IPCs and oRGs, a feature thought to underlie the development and evolution of the large and folded neocortex in higher mammals, including humans (Lui et al., 2011; Borrell and Goetz, 2014; Florio and Huttner, 2014; Sun and Hevner, 2014; Dehay et al., 2015). Consistent with this, exposure to bacterial debris induced folding in the otherwise smooth murine neocortex. This emphasizes that bacterial components could have profound effects on features of human cortical development.

Activation of TLRs in the brain by pathogens is well-known to induce an innate immune response. During bacterial meningitis, PAMPs interact with TLR2/1 to initiate inflammation and neuronal damage (Braun et al., 1999; Tuomanen et al., 1985; Yoshimura et al.,1999). TLRs are expressed in the fetus as early as E10 and activation of TLR2/1 by injecting its agonist (PAM3CSK4) into the embryonic brain ventricle at E15 represses NPC proliferation through a classical inflammatory response (Kaul et al., 2012; Okun et al., 2010; Frost and Schafer, 2016). The interaction of viral RNA and TLR3 represses NPC proliferation by inducing inflammatory signaling via NFkB (Yaddanapudi et al., 2011; Lathia et al., 2008; De Miranda et al., 2010). Neurogenesis in the adult hippocampus is decreased by the absence of TLR2 and increased by loss of TLR4 (Rolls et al., 2007; Monje et al., 2003). However, whether PAMPs might act as morphogens in mammalian neurodevelopment independent of inflammation, as they do in Drosophila (Lemaitre et al., 1996; Morisato and Anderson, 1995), remains unclear. Our data show that activation of TLRs 2 and 6, but not TLR1, increased fetal neurogenesis and permanently altered brain morphology in the absence of inflammation. Thus, by switching partners in the heterodimer at different time points of development, TLR2 participates in either fetal neurogenesis (TLR2/6) or postnatal inflammation (TLR2/1) in mammals.

To explain how bacterial components expanded NPCs, we reasoned that a TLR2/6 developmental pathway must terminate in a nuclear component. FOXG1 is a transcription factor known to promote NPC proliferation by inhibiting FOXO (Seoane et al., 2004). Dysregulation of FOXG1 in humans is associated with changes in brain size and neuropsychiatric disorders (Kortüm et al., 2011; Mariani et al., 2015; Hazlett et al., 2017; al Haddad et al., 2019), a phenotype we replicated in our mouse model prenatally challenged with bacterial infection and treatment. Consistent with these findings, bacterial challenge strongly increased the levels of FOXG1 and decreased FOXO in the embryonic brain. Moreover, bacterial challenge activated PI3K/AKT signaling, which is well known to occur downstream of TLR2 (Pourrajab, et al., 2015) and induces proteosomal degradation of FOXO (Aoki et al., 2004). These changes required TLR2/6, providing a molecular mechanism for PAMP-induced NPC proliferation downstream of TLR2/6.

Surprisingly, spatial transcriptomics revealed an enrichment in genes associated with HH signaling, which has not been previously connected to TLRs. Our data suggest that bacterial debris activated HH signaling indirectly via suppressing cilia and GLI3R formation. Supporting this, a previous study showed that TLR2 promotes disassembly of cilia in cultured cells (Kiseleva et al., 2019). We found that TLR2 localized to the base of cilia in vRGs and that bacterial challenge decreased the number of vRGs with cilia. Thus, TLR2 appears to signal to disassemble the primary cilium at its base. Since vRGs form the wall of the cerebral ventricle and project cilia into the cerebrospinal fluid, the association of TLRs with cilia may promote their function as sentinels. Moreover, as shown for the genetic ablation of cilia in vRGs at early stages of corticogenesis (Wilson et al., 2012; Youn and Han, 2017), the decrease in ciliated vRGs in PAMP-challenged embryos was accompanied by decreased GLI3R levels and derepression of HH target genes, including *Foxg1* and *Fgf15,* which shorten the cell cycle and promote proliferation of vRGs. Importantly, the genetic removal of cilia or GLI3 by E12.5 using a *Nestin::Cre* [Tg(Nes-cre)1Wmz] line but not by E14.5 using a different *Nestin::Cre* [Tg(Nes-cre)1Kln] line expands NPCs and the cortex (Wilson et al., 2012; Wang et al., 2011; Tong et al., 2014). These findings highlight the importance of developmental timing in NPCs’ responses to regulatory factors and provide a possible explanation for the time-dependent effects of bacterial challenges.

We therefore propose a novel neurodevelopmental pathway that drives PAMP-induced NPC proliferation by connecting TLR2/6 to primary cilia and thus to HH signaling and its target genes *Foxg1* and *Fgf15*. The TLR2/cilia/HH/FGF15/FOXG1 and TLR2/PI3K/AKT/FOXO pathways may function in parallel as well as by crosstalk to induce NPC proliferation in the bacterial challenge model. This process is independent of TLR2/1-induced inflammation and neuronal death that characterizes PAMP-induced responses from the innate immune system after birth. Importantly, in control embryos without pathogenic bacterial challenge, the loss of TLR2 or TLR6 increased the basal number of neurons. These findings establish that an endogenous TLR signal regulates normal neurodevelopment in the embryo and that BCW modulates that signal.

---

**Supplementary Figure S1. Timing of expansion of each layer of cortical neurons. Related to Fig 3C**

Sections of E16 fetal brains challenged at E10 as per Fig 3C and injected with EdU (green) on day as indicated. Sections were costained for cortical markers TBR1 (red) or SATB2 (purple). Each image is representative of 2-3 embryos/dam per condition for 3 independent experiments. Arrow: an example of weakly EdU+ cells that were born after multiple divisions following EdU incorporation at E12. Arrowhead: an example of strongly EdU+ cells that were born from a division directly following EdU incorporation at E12.

**Supplementary Figure S2. Effect of loss of TLR signaling on proliferation of neuronal subtypes. Related to Fig 6A**

Mothers deficient in *Tlr1, 2, or 6* vs wild-type were challenged at E10 and fetal brains were harvested at E16. Sections were stained for cortical neuronal markers (TBR1, CTIP2, SATB2) and enumerated. (**O**)= Amp; (•) = Spn + Amp. Each condition was tested in 2-6 embryos from 3-4 independent experiments. Data are shown as mean ± SEM: *P*-values determined by unpaired two-tailed t-test are shown above bars. Each condition was tested in 2-8 embryos from 2-4 independent experiments. Data are shown as mean ± SEM: *P*-values determined by unpaired, two-tailed t-test are shown above bars.

**Supplementary Figure S3. Impact of PAMP challenge on VZ gene expression analyzed by spatial transcriptomics. Related to Fig 7A**

**A)** Dotplot visualizing expression patterns across selected genes of interest, particularly genes known to act downstream of hedgehog signaling. Dot size corresponds to the proportion of capture areas expressing a gene. Color corresponds to the scaled average expression. VZ = ventricular zone. **B)** Mapping of the area of the fetal brain for spatial transcriptomic analysis was determined by manual delineation of anatomical markers for VZ and utilizing spatial transcriptomics to verify region that expressed VZ markers.

**Supplementary Table. Primers for identification of *Tlr* knockout mice. Related to animal methods**

## STAR Methods

### Key Resources Table

#### Resource Availability

Lead contact: Further information and requests for resources and reagents should be directed to and will be fulfilled by the lead contact, Elaine Tuomanen

##### Materials availability

This study did not generate new unique reagents.

##### Data availability

- The data reported in this paper will be shared by the lead contacts upon request.
- This paper does not report original code.
- Any additional information required to reanalyze the data reported in this paper is available from the lead contact upon request.

#### Experimental model and subject details

##### Streptococcus pneumoniae live infection maternal challenge model

Wild type C57BL/6 and *Tlr1^-/-^* mice were obtained from The Jackson Laboratory; C57BL/6 *Tlr6^-/-^* were obtained from Oriental Bioservices Inc.; C57BL/6 *Tlr2^-/-^* (Yoshimura et al, 1999) were bred in house. All mice were maintained in the St. Jude Animal Resource Center BSL1 and 2 housing, and all experiments were carried out in accordance with institutional guidelines and NIH protocols. For maintenance of the colonies and identification of potential genotypes, PCR was performed using the designated primers (Supplemental Table1).

Based on our previous protocol (Humann et al., 2016), timed pregnancies were dated by ultrasound and each experimental group contained 3-5 dams expecting 3-8 pups per dam. Dams were challenged intratracheally at E10 or E11 with 1x10^6^ CFU *S. pneumoniae* strain TIGR4X. Twenty-four hours after challenge, blood titers were obtained to assure consistent bacterial load (mean 1x10^5^ CFU/ml) and the dams were treated with ampicillin (100mg/kg) intraperitoneally twice daily until embryos were harvested for morphological or biochemical analysis. Experiments were repeated a minimum of 3 times.

For some experiments directed at labeling of proliferating cells *in vivo*, bromodeoxyuridine (BrdU 50µg/g, Sigma B5002) or 5-chloro-2’-deoxyuridine (CldU 43µg/g, Sigma C6891), and/or 5-ethynyl-2’-deoxyuridine (EdU 10µg/g, Invitrogen A10044) were injected intraperitoneally into the pregnant dams at designated times before euthanasia for harvest of embryos.

##### Neuronal precursor cell culture

Embryonic brains were harvested at E12.5-E13 in cold PBS and the cortical hemispheres were isolated and separated from the ganglionic eminences and the meninges (Currle et al., 2007). The tissue was then processed using the Brainbits protocol and cells were plated in 12 well dishes at a density of 400,000 cells/ well on polyD-lysine coated plates in complete Neurobasal Media. 24 hours after seeding, cells were washed and pretreated with TLR 2/6 antagonist GIT27 10µg/mL (Tocris 3270) or TLR 2/1 antagonist CU-CPT22 5 µM (Tocris 4884) for 2 hours. Cultures were challenged with purified cell wall (CW) prepared from *S. pneumoniae* as described previously (Humann et al., 2016). Briefly, CW was prepared by dilution in NPC culture media (source catalog #) and sonicated in a cool water bath for 30 minutes before addition to NPCs at a bacterial equivalent of 1x10^4^ CFU. Two hours post-CW treatment, NPCs were lysed on ice for 5 min in cold RIPA buffer (Sigma R0278) with protease inhibitors (CST, 5872S). Cell lysates were collected and centrifuged to collect debris. To separate nuclear and cytoplasmic fractions of treated NPCs, cells were rinsed with cold dPBS, scraped from the plate into freshly made ice cold 100µL fractionation buffer (20mM Hepes, 10mM KCl, 2mM MgCl_2_, 1mM EDTA, 1mM EGTA, 1mM DTT. After 15 min on ice, the cells were passed 10x through a 27 GA needle and incubated another 20 min on ice. Samples were spun 5 min at 3000 RPM and supernatant containing cytoplasm was harvested. The pellet containing the nuclear fraction was resuspended in 100µL fractionation buffer.

#### Method details

##### Immunohistochemistry

###### Frozen sections

Embryonic brains were fixed in 4% paraformaldehyde in PBS overnight, rinsed once in PBS, and cryoprotected in 30% sucrose in PBS at 4°C. Cryoprotected brains were then cut along the midsagittal line and embedded in M1 embedding matrix (ThermoScientific 1310). Tissue blocks were cryosectioned at a thickness of 12 µm and sections were placed on glass slides to dry before storing at - 80°C. For staining, sections were brought to room temperature and placed in boiling antigen retrieval buffer (10 mM sodium citrate, 0.05% Tween 20, pH 6.0) for 10 min. They were briefly rinsed in PBS and placed in blocking solution (5% donkey serum, 0.3% TritonX100 in PBS) for 45 min. Primary antibodies were diluted with 0.1% TritonX100 and applied to the sections and left overnight at 4°C. Sections were rinsed 3 times in PBS and placed in secondary antibody solution at room temperature for 2 hrs. Sections were rinsed 3X in PBS and 2-5µg/ml of Hoechst dye was applied for 10 min at room temperature and rinsed with PBS. Stained sections were mounted in Prolong diamond antifade mounting media (Invitrogen P36961) and stored at room temperature in the dark for 3-4 days to dry.

###### Paraffin sections

Embryonic brains were harvested as above and placed in 10% formalin solution and submitted to the St Jude Veterinary Pathology Core where they were cut along the mid sagittal line or the coronal plane (for cortical folding analysis) and embedded in paraffin and sectioned as 4 µm thick sections. Slides with paraffin sections were stored at room temperature. Slides were first deparaffinized in three rounds of xylene treatment for 5 min each, followed by three 100% ethanol washes for 4 mins. They were washed in 95% ethanol for 4 min followed by 2-3 rinses in distilled water. Sections were transferred to prewarmed antigen retrieval buffer placed in a rice cooker for 15 min. They were then washed in distilled water and PBS and fluorescence staining procedure was conducted as described above for the frozen sections.

###### Staining procedures

Nissl staining was performed with cresyl violet (Sigma C5042) according to the manufacturer’s instructions. EdU staining was performed using the Click-It EdU Alexa Fluor®488 kit (Invitrogen C10632) in accordance with manufacturer’s instructions. For BrdU co-staining, primary and secondary antibody treatment was performed first followed by denaturing in 2N HCl for 30 min at 37°C. Sections were rinsed in 0.1M borate solution (pH 8.5) 3 times for 5 min each followed by PBS washes. Primary antibody solutions containing BrdU or CldU were applied to the sections and placed in 4°C overnight followed by the fluorescence staining method described above. Tyramide Super-BoostTM kit (Invitrogen B40922) was used for triple staining for TLR2, gamma tubulin, and ARL13B following the manufacture’s protocol.

Sections were stained with anti-TBR1 (Abcam ab31940, 1:300), anti-CTIP2 (Abcam ab18465, 1:300), anti-SATB2 (Abcam ab51502, 1:300), anti-TBR2 (eBioscience 14-4875-82, 1:300), anti-PAX6 (R&D AF8150, 1:100), anti-Ki67 (Abcam 15580, 1:100), anti-CldU (NovusBio NB500-169, 1:100), anti-gamma tubulin (Sigma Aldrich T5192, 1:1000), anti-ARL13B (UC Davis NeuroMap, 1:2000), anti-TLR2 (Abcam ab209216, 1:100), DAPI (Sigma Aldrich D9542, 1ug/mL), or Hoechst 33342 (Biotium 40046, 2-5ug/ml). The following secondary antibodies were used at a dilution of 1:500 and incubated at room temperature (20-23 °C) for 2 h: CF^®^ 488A-conjugated donkey anti-rat IgG (Biotium 20027), donkey anti-rabbit IgG (Biotium 20015), donkey anti-mouse IgG (Biotium 20014); CF^®^ 568-conjugated donkey anti-rabbit IgG (Biotium 20098), donkey anti-mouse IgG (Biotium 20105); CF^®^ 647-conjugated donkey anti-rat IgG (Biotium 20843), donkey anti-rabbit IgG (Biotium 20047), donkey anti-mouse IgG (Biotium 20046); Alexa Fluor (AF) 647-conjugated anti-goat (Thermo Fisher Scientific A-21447, 1:200), AF 488-conjugated anti-mouse IgG (Thermo Fisher Scientific A-28175, 1:200), and AF 568-conjugated anti-rabbit (Thermo Fisher Scientific A-11011, 1:200).

#### Microscopy

##### Stereology

Methods for calculations of total neuronal cell populations of the cortex and hippocampus were as described previously (Humann et al., 2016). Briefly, both frozen and paraffin-embedded sections were stained with cresyl violet, and standard stereology methods using StereoInvestigator with Cavalieri estimator and the optical fractionator probe (MBF Biosciences) were implemented. Both right and left sides of the cortex were measured and the average of 3 sections per embryo was calculated. Measurements were taken for 3-5 dams with 4-8 embryos each for each experimental group.

##### Fluorescence image acquisition and analysis

Images were acquired on a Nikon C2 (Nikon NIS elements software) or a 3i Marianas system (Slidebook software) using a 40x or 63x objective. For frozen sections of 12 µm thickness, a *z-*stack of 13 optical sections at a step size of 0.34 µm and a tiling of multiple images were combined for quantification analysis. For paraffin sections of 4 µm thickness, a 2D montage of a tiled image of the cortical plate region of the right hemispheres was captured using a 40X oil objective on the Marianas. For all experiments 1-3 sections from single embryos were imaged. Measurements from 2-3 dams for each experimental group, with up to 3-7 embryos each were quantified. Quantifications of neuronal number were carried out using Imaris x64(Bitplane) image analysis software.

##### Cell cycle analysis

We injected CldU at 2 h before harvest and EdU at 0.5 h before harvest (Hayes and Nowakowski, 2000). EdU+ PAX6+ TBR2-cells represent vRGs in S phase (S-cells) at the time of harvest, whereas CldU+ EdU-cells represent vRGs that have left S phase (L-cells) and entered G2 phase during the 1.5 h between CldU and EdU injections. The ratio of the length of a period of the cell cycle to that of another period is equal to the ratio of the number of cells in each period (Nowakowski et al., 1989). Thus, we determined the length of S phase (Ts) from the equation 1.5 h/Ts = L-cells/S-cells, then the length of cell cycle (Tc) from the equation Ts/Tc = S-cells/Proliferating RGs (Ki67+PAX6+TBR2-; Ki67 labels all phases of cell cycle).

##### Cortical folding analysis

Nissl stained coronal sections of P10 harvested brains were imaged with the Axioscan at 20x and stitch-fused using Zen Blue. Folds in the cortex were counted from 2 dams per experimental group with 2-8 embryos each.

### Embryonic brain lysis

Individual embryonic brain cortices challenged *in utero* were dissected just after harvest and lysed in 250µL ice cold RIPA buffer (Sigma R0278) with protease inhibitors (CST 5872S). The tissue was homogenized and vortexed for 2 hours at 4°C. Lysates were centrifuged to collect debris and supernatant was harvested.

### Western blot analysis

Concentration of total protein in each sample was determined by BCA (Thermo Scientific), and equal amounts were loaded on a 4-12% SDS PAGE. Antibodies used for western blot analysis included: anti-FOXG1 (Abcam ab18259, 1:1000), anti-FOXO1 (CST 2880, 1:1000), anti-GLI3 (R&D Systems AF3690, 1:200), anti-PI3K (Thermo PA5-19833, 1:1000) anti-phospho-AKT (CST 4060, 1:1000) anti-AKT (CST 4691, 1:1000), anti-beta-actin (Sigma A5441, 1:20,000)

### RNA isolation and quantitative real time PCR

Individual embryonic brain cortices (cortical plate regions with VZ, devoid of meninges and ganglionic eminences) were harvested, snap frozen and stored at -80°C for RNA extraction. RNA was extracted using the RNAeasy kit (Qiagen 74104), according to manufacturer’s instructions. cDNA was synthesized from purified RNA with the Superscript III First-Strand Synthesis kit (Invitrogen 18080-400) according to manufacturer’s protocol. RT-PCR was run using Taqman Fast Advanced Master Mix (Applied Biosystems 4444557) and custom TaqMan Gene Expression Assays *FoxG1* Mm02059886_s1, *Fgf15* Mm00433278_m1 and endogenous control *Gapdh* Mm99999915_g1 (Applied Biosystems). 3-4 dams per condition with 5-8 samples were analyzed and each sample was run in triplicate reactions.

### Behavioral analysis

Pups born to mothers treated with Spn and ampicillin or ampicillin alone at E10 were assessed for proper cognitive function, social and repetitive behaviors at 2, 4 and 6 months of age. To test working memory, the spatial recognition test was used as previously described (Humann et al., 2016; Bevins and Besheer, 2006; Crawley, 2007). Briefly, mice were exposed to 2 of 3 arms in a Y-maze for 8 min with the third arm gated. One hour after returning to its cage, the mouse was allowed to explore all 3 arms for 5 min. Time and entries were noted for all arms. To assess repetitive behavior, mice were video recorded alone in their cage for 5 minutes and the time spent grooming was calculated. To study sociability, a white box divided by a perforated plexiglass was used. Experimental mice were acclimated to one side of the box for 8 min with the other side empty. The mouse was returned to its cage for one hour, then subsequently, placed back in the box with a non-littermate mouse on the other side of the divide. The number of approaches to the divider as well as time spent attempting to interact with the non-littermate was calculated. All behavioral assays were recorded by video and double checked for accuracy.

### Spatial transcriptomics

Embryos were obtained at E12, and whole heads were dissected out for analysis with the Visium Spatial Transcriptomics kit (10X Genomics PN-1000184). Immediately after dissection, samples were frozen in isopentane chilled by liquid nitrogen, embedded in OCT embedding matrix (Sakura 4583), and then stored at -80°C. 10 µm sagittal sections were obtained following 10X Genomics suggested practices. Tissue permeabilization was optimized by testing 0, 3, 6, 12, 18, 24, and 30 minute incubations on test sections, with 12 minutes determined as optimal. A single representative section was then obtained from each of four mice per condition, and two sections were placed on each Visium slide (i.e., one section from each condition). Sections were imaged on a Nikon Eclipse N*i*-E scope, and slide fiducials and tissue coverage areas were manually annotated using Loupe Browser (10X Genomics). Libraries were prepared according to manufacturer recommendations and sequenced on an Illumina NovaSeq platform with a sequencing configuration of 28-10-10-120 (R1-i7-i5-R2) at >200 million clusters per library. Data were analyzed using SpaceRanger (v1.2.1, 10X Genomics) using the corresponding mm10 reference, with the reverse read trimmed to 90 base pairs.

Downstream analyses were performed using the STUtility (v0.1.0; Bergenstråhle et al., 2020) and Seurat (v4.0.3; Hao et al., 2021) packages in the R environment (v4.1.0), with each library independently processed using SCTransform (regressing out the percentage of mitochondrial expression per spot) and then merged into a single object. Capture areas overlaying putative ventricular zone regions were identified by anatomical recognition based on the prenatal mouse brain atlas (Schambra, 2008) and confirmed by the abundance of *foxg1* expression in these sites., and analyses focused on these regions. Differentially expressed genes (DEG) between conditions were determined using the Seurat FindMarkers function with default parameters, with *P*-values adjusted for multiple comparisons using Bonferroni correction. Gene Set Enrichment Analysis (GSEA; Subramanian et al., 2005) was performed using GSEAPreranked on a DEG list obtained using FindMarkers with min.pct set to 0.01 and logfc.threshold set to 0, with ranks ordered by average log2 fold-change; for this analysis, Hallmarks (v7.4; Liberzon et al., 2015) gene sets were queried using the Mouse MSigDB symbol remapping chip (v7.0) with a classic enrichment statistic.

### Statistics

Detailed statistical parameters are found in each figure legend.

All ***in vitro* experime**nts were conducted in at least three independent biological replicates. Positive and negative controls ensure unbiased data interpretation.

***In vivo* experiments Sex:** Female pregnant mice began each experiment; All male and female embryos were analysed. Group Sizes: Groups contained least 2-3 pregnant mice that then yielded at least 4-6 embryos/dam, a cohort size giving a power of 0.95 to detect a 0.5 log change in cell number at a P-value of 0.01. The treatment assignment, challenge status and endpoint analysis was blinded to investigators. Data Analysis to achieve robust and unbiased results: Unless otherwise specified, comparisons of phenotypes were analyzed by calculating the mean and standard deviation. P values will be calculated using a two-tailed, unpaired Student’s t-test for two-group comparisons or using two-tailed T test with Welch’s correction or a Mann-Whitney test in Prism version 6.05 (GraphPad). In instances where multiple experimental conditions are compared with a single control group, statistical significance is tested using one-way analysis of variance (ANOVA) followed by Bonferroni’s multiple comparison postcomparison test. A P value less than 0.05 was considered significant.

#### Analysis of brain slices

For consistent quantification of progenitor cells in slices, the VZ was defined as the area lining the ventricle and containing dense PAX6+ TBR2− nuclei up to the area where cells uniformly express TBR2 (RGs and newborn IPs express PAX6; IPs express TBR2); the SVZ as the second cell-dense area containing uniformly TBR2^+^ cells above the VZ and below a cell-sparse area. Continuous stretches of TBR2+ cells form the boundary between the VZ and SVZ. We defined oRGs as PAX6+ TBR2− cells above the VZ. Counting was done by at least 2 blinded individuals to ensure unbiased analysis. We used right and left hemispheres of all embryos from at least 3 dams per experimental group and repeated all experiments at least 3 times to ensure reproducibility.

#### Spatial transcriptomics

Data was generated and analyzed under the direction of Dr. Jeremy Crawford who was blinded to the experimental groups. Spatial transcriptomics data was generated using standard protocols including the 10X Genomics Visium approach. Library sequencing was performed at our institutional core facility using the Illumina NovaSeq platform. Sequencing data was processed using Space Ranger (spatial) programs (10X genomics), and the resulting expression matrices were analyzed using several open-access analytical tools, including the Seurat R package. Changes in gene expression revealed by spatial transcriptomics approaches were confirmed by measuring transcript abundance via qRT-PCR in independent samples. To ensure that the conclusions drawn from spatial transcriptomics experiments were rigorous, experiments included multiple, biologically independent sample replicates for each point of comparison. All statistical analyses are corrected for Type 1 error (false discovery rate) by adjusting P-values using the Benjamini & Hochberg method. All data and accompanying documentation are archived on servers both in the laboratory and in the St. Jude Children’s Research Hospital cloud which is backed up daily. All raw sequencing data (and corresponding imaging data for spatial transcriptomics) will be made publicly available via the NCBI Short Read Archive upon publication.

## Acknowledgements

The authors would like to thank the St. Jude Animal Resource Center and St. Jude Small Animal Imaging Center, especially Melissa Johnson, RLAT, for help with timed matings and ultrasound data collection and analysis. We thank the St. Jude Cell and Tissue Imaging Center for microscopy assistance (supported by SJCRH and NCI P30 CA021765-35). Dr. Timothy Flerlage was instrumental in directing us to spatial RNA-Seq. This work was supported by R01AI128756 to ET, R01NS100939 to Y-GH, R01AI135025 (sub to PT), U01AI150747 to PT, and ALSAC.

## Author contributions

Conceptualization, E.I.T. and Y-G.H.; Methodology, K.R., J.L., B.M., G.G., Y-H.Y., C.G., C-H.C., D.D., S.T., J.C.C.; Investigation, K.R., J.L., B.M., A.J.L., G.G., Y-H.Y. and J.C.C.; Writing – Original Draft, B.M., J.C.C., E.I.T. and Y-G.H.; Writing – Review & Editing, B.M., J.C.C., P.T., Y-G.H. and E.I.T.; Funding Acquisition, E.I.T., P.T., and Y-G.H.; Resources, E.I.T., P.T., and Y-G.H.; Supervision, E.I.T. and Y-G.H.

