## Supplementary Figures and Table for "Fetal neural progenitors process TLR signals from bacterial components to enhance proliferation and rework brain development": Supplementary materials.pdf

### Supplementary Figure S1

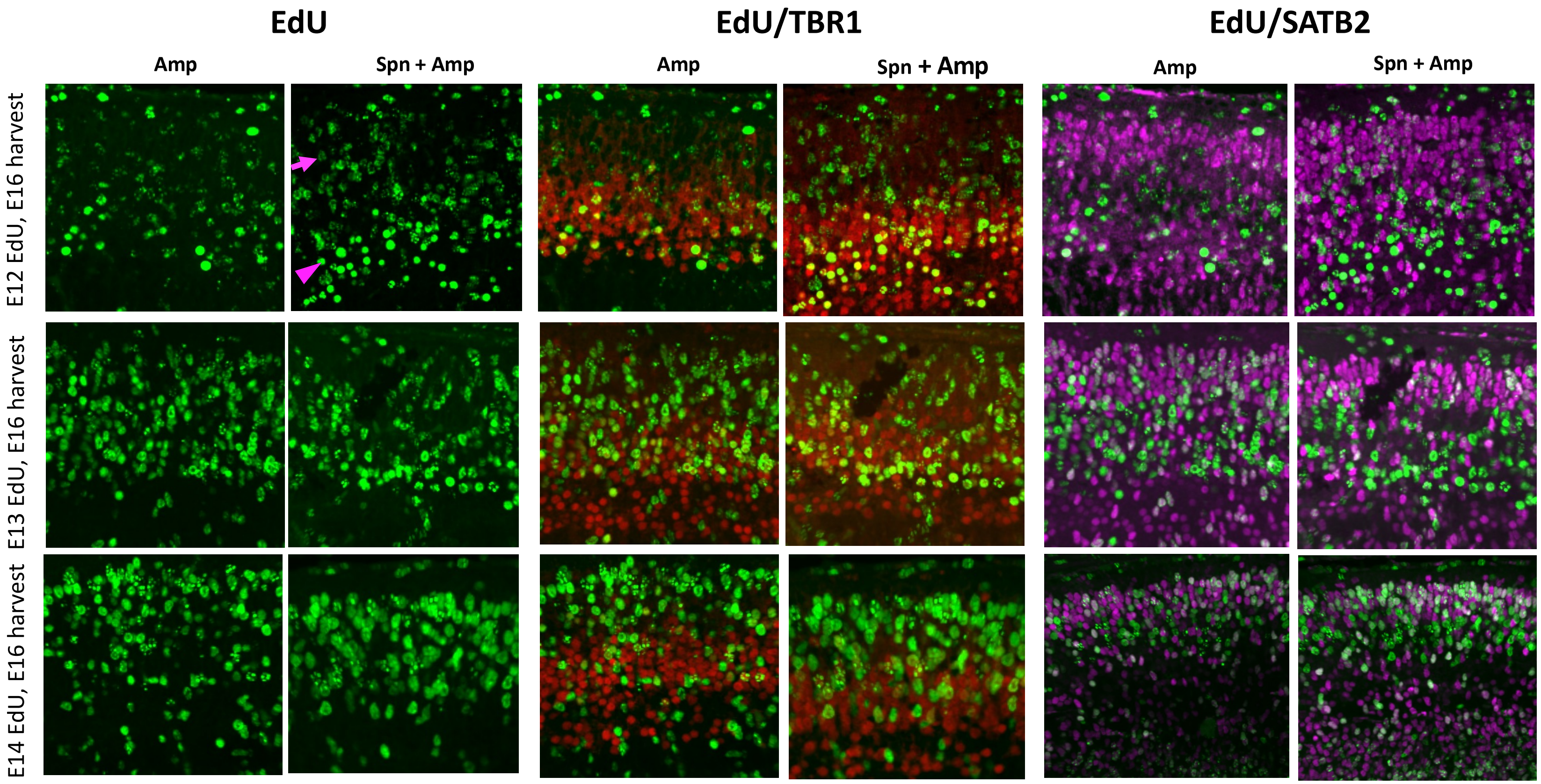

### Supplementary Figure S2

#### Cortical Neurons

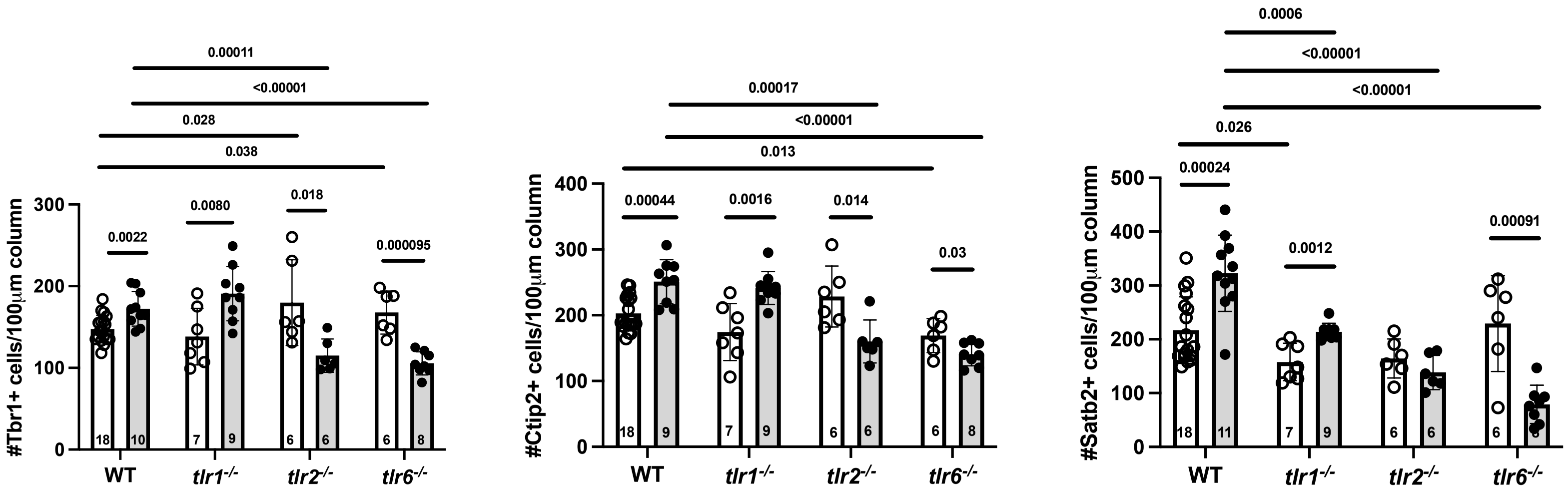

### Supplementary Figure S3

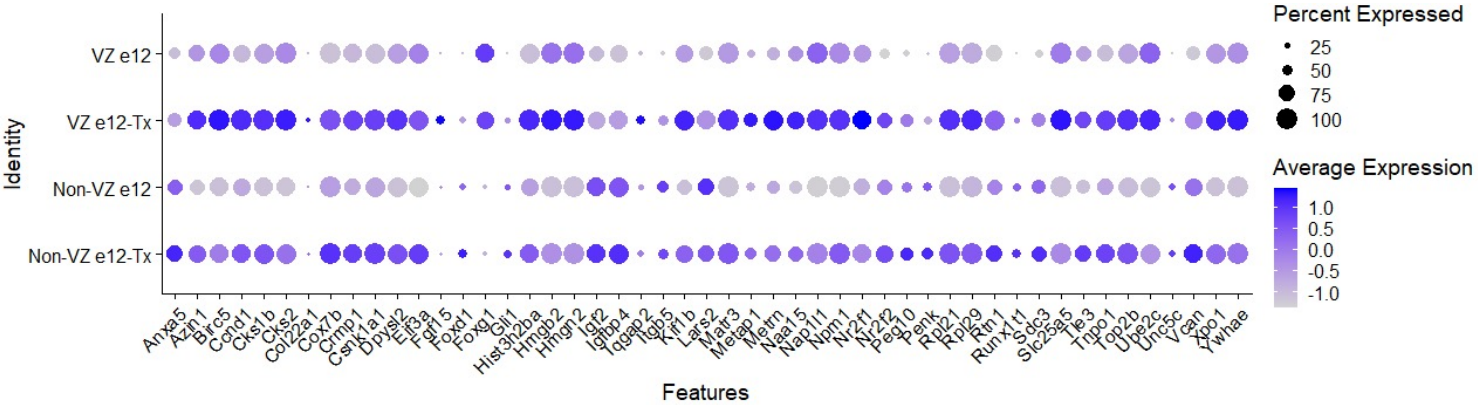


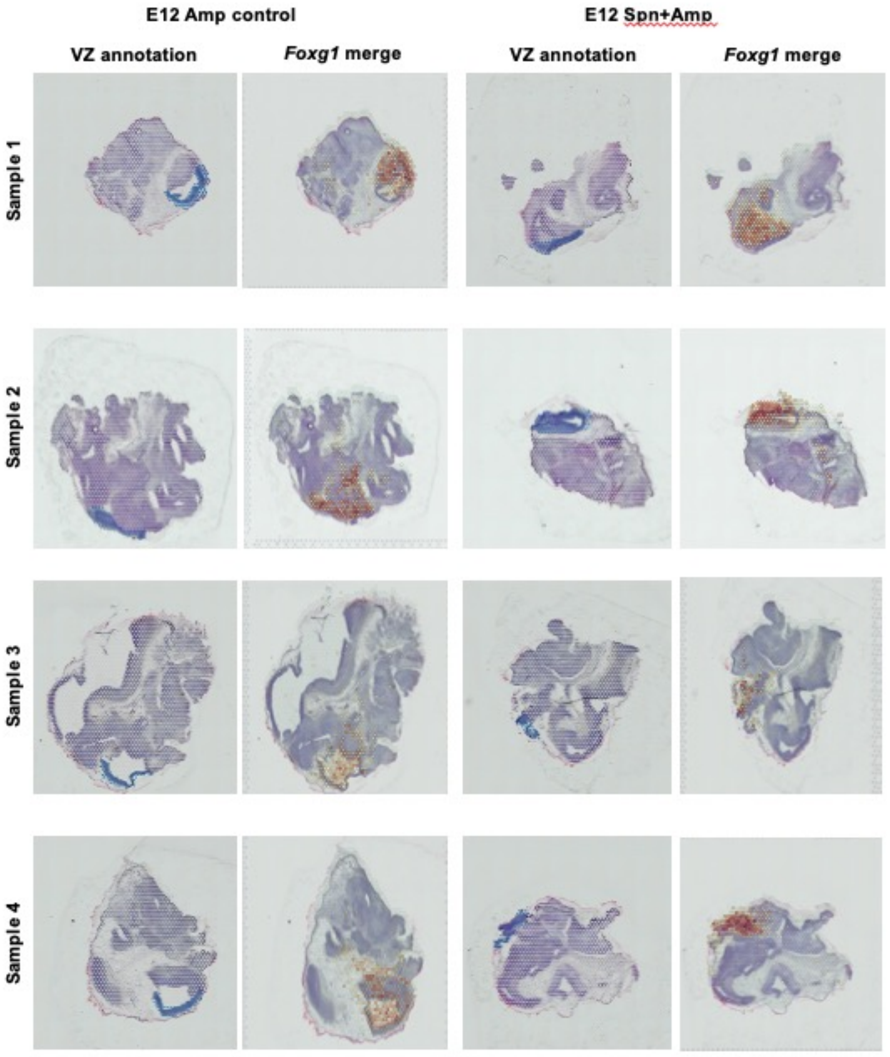

**Supplementary Table 1. Primers for identifying knockout mice.**

| Strains | Vendor | Primers Name | Sequence 5'-3' |
|---|---|---|---|
| *Tlr1 KO* (B6.129S1_*Tlr1*) | Jackson | oIMR6966 | GCCAAACGCAAA CCTTACCAGAGTG |
| | | oIMR6967 | ACGGACACATCCAGAAGAAAACGG |
| | | oIMR6968 | TTCGGCTATGACTGGGCACAACAG |
| | | oIMR6969 | TACTTTCTCGGCAGGAGCAAGGTG |
| *Tlr2* KO | SJCRH | PM.STJ.583 | CTTCCTGAATTTGTCCAGTACA |
| | | PM.STJ.584 | GGGCCAGCTCATTCCTCCCAC |
| | | PM.STJ.585 | ACGAGCAAGATCAACAGGAGA |
| *Tlr6* KO | Oriental Bioservices, Inc. | Wild | GAAATGTAAATGAGCTTGGGGATGGCG |
| | | Extra | TAATCAGAACTCACCAGAGGTCCAACC |
| | | Neo | ATCGCCTTCTATCGCCTTCTTGACGAG |
